# Mutation detection in candidate genes for parauberculosis resistance in sheep

**DOI:** 10.1101/014035

**Authors:** Bianca Moioli, Luigi De Grossi, Roberto Steri, Silvia D’Andrea, Fabio Pilla

## Abstract

The marker-assisted selection exploits anonymous genetic markers that have been associated with measurable differences on complex traits; because it is based on the Linkage Disequilibrium between the polymorphic markers and the polymorphisms which code for the trait, its success is limited to the population in which the association has been assessed. The identification of the gene with effect on the target and the detection of the functional mutations will allow selection in independent populations, while encouraging studies on gene expression. The results of a genome-wide scan performed with the Illumina Ovine SNP50K Beadchip, on 100 sheep, 50 of which positive at paratuberculosis serological assessment, identified two candidate genes of immunity response, the PCP4 and the CD109, located in proximity of the markers with different allele frequency in positive and negative sheep. The coding region of the two genes was directly sequenced: three missense mutations were detected: two in the PCP4 gene and one in the second exon of the CD109 gene. The PCP4 mutations had a very low frequency (.12 and .07) so making hazardous to hypothesize their direct effect on immune response. On the contrary, the mutation detected in the CD109 gene showed a strong linkage disequilibrium with the anonymous marker. Direct sequencing of the DNA of sheep of different populations showed that disequilibrium was maintained. Allele frequency at the hypothesized marker associated to immune response, calculated for other breeds of sheep, showed that the marker allele potentially associated to disease resistance is more frequent in the local breeds and in breeds that have not been submitted to selection programs.

## INTRODUCTION

The aim of marker-assisted selection (MAS) is to select specific DNA variations that have been associated with a measurable difference on complex traits. It is based on the Linkage Disequilibrium (LD) between the polymorphic markers and the polymorphisms which code for the trait; the marker must by necessity be close to the functional mutation for sufficient population-wide LD between the marker and the gene. Also for this reason, success of MAS has been limited in livestock and few studies reported substantial gains with real populations (Dekkers 2004). The possibility to detect the direct markers (i.e. polymorphisms that code for the functional mutations) opens the way to wider fields of applications, on one side because selection could be performed in independent populations, with no care of the LD extent; on the other side, the identification of the gene having major effect on the trait will allow a deeper knowledge of the function and the expression of the gene, as well as of the related gene network, and such knowlege might be of use also to other species and different target traits.

Paratuberculosis, or Johne’s, disease is a chronic granulomatous enteritis, which affects ruminants, caused by Mycobacterium avium subsp. Paratuberculosis (MAP). It causes substantial reduced production, higher susceptibility to acquire other diseases and infertility (Kennedy and Benedictus 2001). In a previous work (Moioli *et al*. 2015), a Genome Wide Association Analysis (GWAS) with the Ovine Bead chip 54K was performed on two groups of ewes that had been found either positive or negative when serologically screened for paratuberculosis diagnosis with the serum antibody ELISA test. These authors hypothesized the presence of 30 putative candidate genes of disease susceptibility, these genes being in close proximity of the polymorphic markers which showed a significant effect on the obtained serological value. While the majority of these genes did not appear to have a straightforward and recognizable effect on disease resistance, but they were either activators of transcription, growth factor activators, or responsible for basic cellular processes, two of them, the Purkinje Cell Protein 4 (PCP4), and the CD109 genes appeared worth to be studied in more depth for their potential role in influencing the immune system. In fact, Jacobson *et al*. (2009), in the murine B cell co-receptor complex, demonstrated an increased expression of PCP4, in wild type mice compared to complement-deficient animals, and correlated the increased expression of PCP4 with B cell maturation into end stage phenotypes. On the other hand, the CD109 belongs to the complement gene family and is a co-receptor of the Transforming growth factor beta (TGF-β), a type of cytokine which plays a role in immunity (Hashimoto *et al*. 2009).

The present study was conceived to search for the presence of further polymorphisms in the coding or regulatory region of PCP4 and CD109 genes, that might explain and corroborate the effect of the two anonymous markers identified by Moioli *et al*. (2015) in influencing disease resistance.

## MATERIALS AND METHODS

Genotyping results at the Ovine 54K Bead chip for 100 sheep of the Sarda breed were available from the previous study (Moioli *et al*. 2015); half of the sheep had been assessed as serologically positive to paratuberculosis, according to the ELISA test; the remaining half were negative, the kit for diagnose considering positivity S/P values > 70. DNA of 42 of these sheep, equally distributed between positive and negative was also made available by the Animal Health Intitute of Viterbo, Italy, that had performed the MAP diagnose.

The genomic scaffolds encoding the PCP4 and the CD109 genes were obtained from the NCBI data base (https://www.ncbi.nlm.nih.gov/genome/?term=ovis+aries); the size and the position of the coding regions of the PCP4 and the CD109 genes on the corresponding chromosomes – OAR1 and OAR8 were identified by performing a standard nucleotide blast (Altschul *et al*. 1990) of the genomic scaffolds against the expressed sequence of the gene under investigation in the NCBI (www.ncbi.nlm.nih.gov/nuccore).

Under the assumption that the markers identified by Moioli *et al*. (2015) be in LD with the polymorphism responsible for the trait variation, and because the average distance between markers of the 50K BeadChip ranges from 60K to 200K, the genomic scaffolds encompassing the PCP4 and the CD109 genes were searched for the presence of further markers. From the genotyping results made available by the previous study (Moioli *et al*. 2015), the allele frequency in the two sheep samples – 50 serologically positive and 50 negative respectively - at each of the markers included in such scaffolds were compared using the χ^2^ test of significance of differences. In fact, χ^2^ test values are deemed to provide solid indication of the region in which linkage disequilibrium had been maintained (Lewis 2002).

Primer pairs to amplify the expressed regions of the genes were designed on the ovine sequence by using Primer3Plus software (www.bioinformatics.nl/cgi-bin/primer3plus/primer3plus.cgi). Direct sequencing of the coding regions of the putative candidate genes was performed on the 3500 Genetic Analyzers (Applied Biosystems, Foster City, CA, USA). DNA of the sheep was kindly provided by Istituto Zooprofilattico del Lazio e della Toscana, Viterbo, Italy.

In a first step, DNA of 10 individuals, 5 serologically positive and 5 negative was amplified at each amplicon, under the assumption that this number was sufficient to detect the presence of further SNPs. Once the novel SNP had been detected, direct sequencing of the amplicon was performed on 42 sheep, equally distributed between serologically positive and negative. Allele frequencies and linkage disequilibrium measures, for both the anonymous markers and the novel detected polymorphisms, were estimated using the Allele Procedure in SAS (SAS Inst.Inc., Cary, NC). Because all the sheep assessed for paratuberculosis diagnose belonged to the Sarda breed, to verify whether LD between the novel detected SNPs and the Ovine Bead chip anonymous markers were maintained also in different breeds, DNA of 33 more sheep of two Italian breeds – Altamurana and Comisana – were directly sequenced at the amplicons encompassing the coding regions of the putative candidate genes. These breeds and individuals were chosen because their DNA was available in the laboratory performing the present study, and because their genotyping results at the Ovine 54K Bead chip were also available, thanks to a previous biodiversity project (Ciani *et al*. 2014).

## RESULTS

### PCP4 gene

The Ovine 50K BeadChip contained 4 markers (Table 1) in the genomic region encompassing the PCP4 gene, two of the markers, OAR1_278884883.1 and OAR1_278980576.1, showed significant difference in allele frequency, between serologically positive and negative sheep, after the χ^2^ test results (Table 1). The nucleotide blast (Altschul *et al*. 1990) of the genomic scaffold NW_004080164.1 against the expressed sequence of PCP4 (XM_004003899.1) showed that the gene is composed of 3 exons (Fig. 1) the two markers with significant difference in allele frequency, OAR1_278884883.1 and OAR1_278980576.1, being located respectively in the first intron and 29 K downstream the stop codon of the gene. Therefore, hypothesizing that putative mutations influencing the target trait be located either in the first part, or in the 3’UTR of the gene, two amplicons were designed, the first, of 346 bp, encompassing exon 2 and part of the flanking regions; the second amplicon, 580 bp, including exon 3 and the whole 3’ UTR region (Table 2). Direct sequencing of the amplicons of Table 2 allowed the detection of one missense mutation in exon 2 (XM_004003899.1; g.97 G>C) producing the aa change from Glu to Gln (Table 4). Moreover, five more mutations were detected in the amplicon encoding exon 3 and the 3’ UTR region, one of them was a missense mutation (XM_004003899.1; g.107 G>A) producing the aa change from Gly to Glu. Allele frequency was reported in Table 4.

**Tabel 1.**
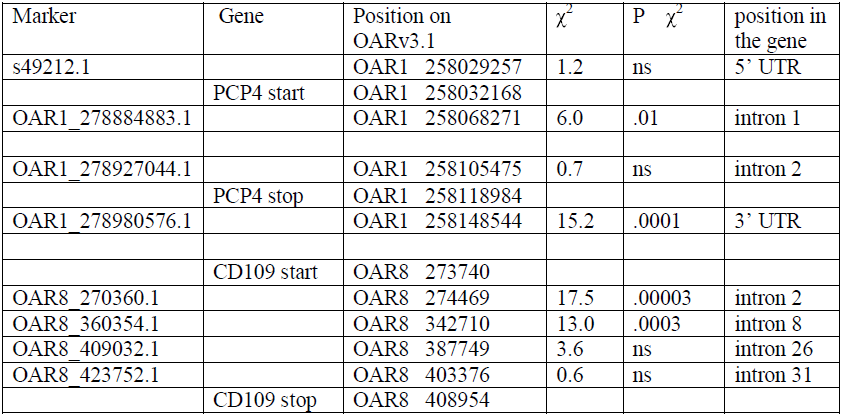
χ test of differences between 50 serologically positive and 362 50 negative sheep at the 363 markers falling in the region around the PCP4 gene (genomic scaffold NW_004080164.1).

**Fig 1.**
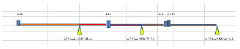
Structure of the PCP4 gene in Ovis aries with indication of the position of the markers of the Illumina Ovine BeadChip

**Tabel 2.**
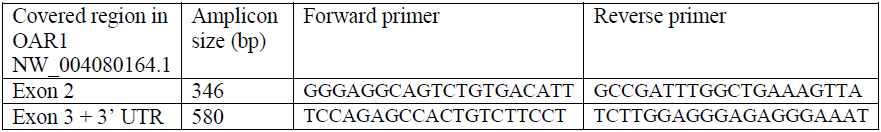
Primers used to amplify the PCP4 gene

**Tabel 3.**
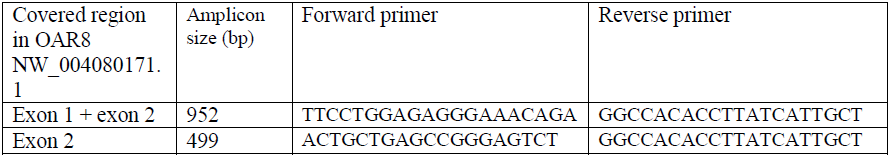
Primers used to amplify the CD109 gene

**Tabel 4.**
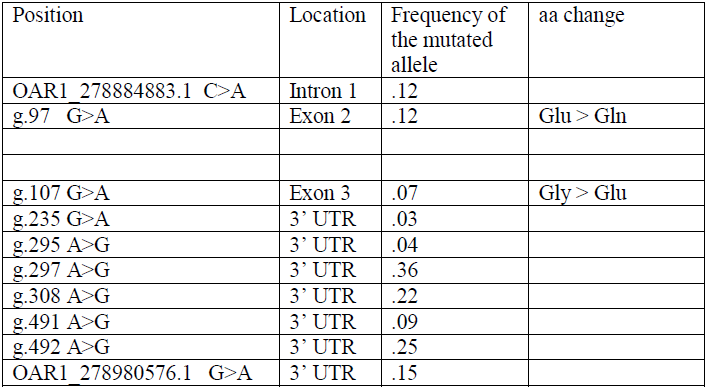
Detected mutations in the 381 PCP4 gene (Acession XM_004003899.1)

The novel detected SNP in exon 2 and the marker in intron 1 (OAR1_278884883.1) showed no LD between each other (.05; Table 5).

**Tabel 5.**
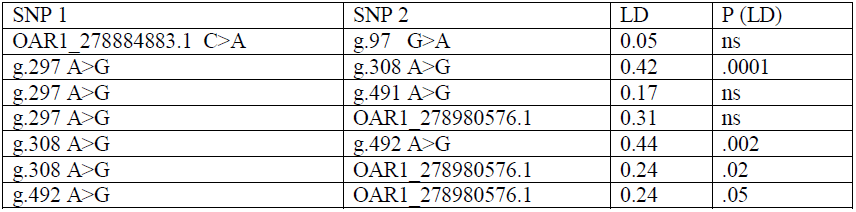
Linkage disequilibium measures between pair of SNPs in the 3’ UTR of the PCP4 gene (Acession XM_004003899.1)

For the novel detected SNPs in the 3’ UTR, LD between pair of SNPs was calculated for the markers with minimum allele frequency (MAF) higher that .10 and reported in Table 5; LD was week (r < .45) for all markers, and significant only in few cases.

### CD109 gene

Table 1 reports the 4 detected markers located in the CD109 gene. The results of the the χ^2^ test of differences between 50 serologically positive and 50 negative sheep showed that only the two markers located in the upstream region of the gene have different frequencies in the two groups.

The nucleotide blast (Altschul *et al*. 1990) of the Ovis aries breed Texel chromosome 8, Oar_v3.1, whole genome shotgun sequence (NCBI Reference Sequence: NC_019465.1) against the expressed sequence of CD109 gene (XM_004011463.1) indicated that the gene encompasses position 20284 to 155500 of the scaffold NC_019465.1 and that the gene is composed of 34 exons (Fig 2). Only two markers, OAR8_270360.1 and OAR8_360354.1 showed significant difference between serologically positive and negative sheep (Table 1), therefore, hypothesizing the presence of putative mutations in LD with the markers, in a first step it was decided to sequence the first exons of the gene. It was moreover noted that 298 nt were missing in scaffold NC_019465.1, i.e. from position 20382 to 20672 nt, so one amplicon was designed in order to identify the missing nucleotides, this amplicon encompassing exon 1, exon 2 and part of the flanking regions (Table 3); direct sequencing of ten individuals allowed the detection of a novel SNP in exon 2 (g.128 A>G; XM_004011463.1). To amplify the whole exon 2, a shorter amplicon was designed, with the forward primer on the novel identified sequence (Table 3).

**Fig 2.**
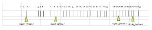
Structure of the CD109 gene in Ovis aries, with indication of the position of the markers of the Illumina Ovine BeadChip

The novel detected mutation in exon 2 had a frequency of .25 in the positive and .54 in the negative sheep (P<0.01), and it was a missense mutation producing the aa change from Glu to Arg. Allele frequency at this mutation was .25 in the positive and .54 in the negative sheep (Table 6). LD between the missense mutation and marker OAR8_270360.1 was significant (LD=0.74; P<.0001).

**Tabel 6.**
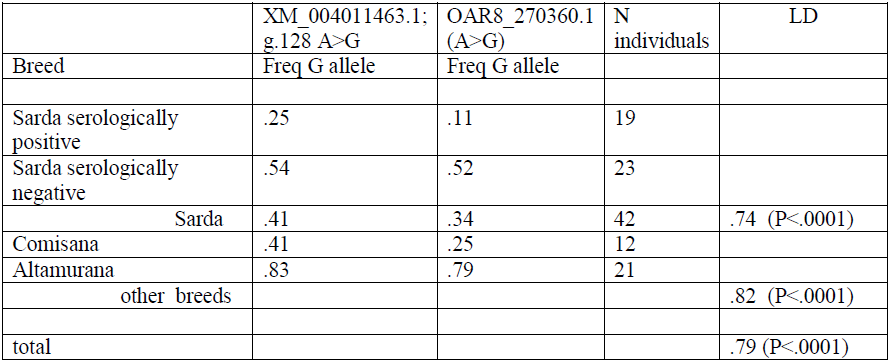
Detected mutations in the CD109 gene and LD measures beween the mutation and the marker.

To verify whether LD between marker OAR8_270360.1 and the novel detected mutation were maintained also in different sheep populations, the amplicon encoding the Glu/Arg mutation was sequenced in 33 more sheep, which had already been genotyped at the Ovine 54K Bead chip in previous projects (Ciani *et al*. 2014; Moioli *et al*. 2013). The conservation of LD between the marker and the missense mutation in breeds different than the Sarda would in fact substantiate the use of this marker, as a marker natural immunity, for any individual which was being genotyped for different purposes – genomic selection or biodiversity studies – with no need to direct sequencing the missense mutation.

Table 6 reports the frequency of the novel detected mutation in exon 2 of the CD109 gene for the Comisana and Altamurana, making evident the different frequencies of the G allele. The two breeds had also opposite allele frequencies at marker OAR8_270360.1 (Table 6). As in the Sarda breed, LD between marker and missense mutation was maintained also in the Comisana and Altamurana breeds: 0.82 (P<.0001).

## DISCUSSION

The present study had been formulated under the assumption that anonymous markers previously associated with disease resistance were in LD with missense mutations in the genes located in proximity of the markers, so to corroborate the influence of the gene on the target trait. In the present study, two putative genes were scanned: PCP4 and CD109. The analysis of the PCP4 gene allowed the detection of a missense mutation in exon 3 and four more SNPs, in the 3’ UTR region; however, the low frequency of the missense mutation and the week LD between the other SNPs and the marker did not endorse a possible direct role of this portion of the gene in influencing the immune system. The discovery of another marker in intron 1 of the PCP4 gene, with different allele frequency between seropositive and negative sheep suggested to perform a further genomic scan in the first part of the gene, which allowed the detection of a further missense mutation in exon 2, this mutation, however, having a very low frequency in the analyzed population (.12). Furthermore, no LD was registered between the mutation and the anonymous marker OAR1_278884883.1. In conclusion, for the PCP4 gene, the suggestive association proposed by Moioli *et al*. (2015) might have been simply probabilistic, due to the limited size of the analyzed sample and the possible relationship between individuals of the same flock.

On the contrary, the detection of a missense mutation in the CD109 gene was intriguing, the OAR8_270360.1 marker, hypothesized as a marker of natural immunity, being located only 137 bp upstream this mutation. But even more striking was the extent of LD between the marker and the mutation (0.79; P<0.0001), which was maintained also in breeds different from the Sarda and in breeds were the mutation had opposite allele frequencies. For the Sarda breed, allele G was the minor allele at marker OAR8_270360.1, with frequency =.34; but the serologically positive Sarda sheep showed a much lower frequency (.11) while the negative sheep had a frequency of .52 (Table 6). Similar trend was detected for the mutation in LD with the marker. In the Comisana breed the G allele frequency was even lower than in the Sarda (.25). Allele G was on the contrary the major allele in the Altamurana breed, this fact encouraging to perform further investigations. Tha Sarda and the Comisana breeds are two specialized Italian dairy breeds of sheep, which have been submitted to breeding programs for the improvement of milk yield, and have rapidly spread outside of their original geographical areas, so that they have largely replaced indigenous and low-productive breeds of sheep all over Italy, like the Altamurana. The latter is a dairy sheep, belonging to the subgroup of south European milk sheep (Pieragostini and Dario 1996), living in harsh environment, and producing no more than 50 litres milk in a short lactation. Because the Altamurana breed, as well as several other Italian indigenous breeds, is maintained for conservation purposes, the discovery of markers in genes which play a role in the immune systems, with opposite allele frequency in selected and indigenous hardy breeds, would corroborate the usefulness of developing conservation programs for the endangered breeds. The exam of the genotyping results of 992 sheep of 20 breeds, obtained from previous genome-wide analyses of genetic diversity (Ciani *et al*. 2014), results which were kindly made available by the authors for improving the present study, showed that the average frequency of the G allele at marker OAR8_270360.1, i.e. the hypothesized allele associated to the trait “disease resistance”, was .44, with the majority of the breeds with frequency ranging between .40 and .50. The lowest frequency was obtained by the Sarda (.35), Comisana (.31), Delle Langhe (.27) and Gentile (.25); the first three breeds are the only Italian breeds under selection for the improvement of milk yield; the Gentile is the Italian merino, voluntarily improved during the past centuries with the use of foreign merino rams. On the other hand, the highest frequency of the G allele was obtained in the Massese (.56), Fabrianese (.61) and Istrian (.67). The latter is particularly interesting because the study by Ciani et al. (2014) highlighted a genetic closeness of this breed with other breeds of Northern Italy (Bergamasca, Biellese and Sambucana) where the frequency of the G allele was among the lowest (.31-.43). Istrian sheep were bred mainly for their unusual characteristics: their distinct long-stepping walk and ability to graze in rocky terrain (Kompan *et al*. 1999). Also the Massese is an indigenous hardy dairy breed, grazing all the year in the hills and mounatins of Tuscany (Casoli *et al*. 1989). The Fabrianese is a course wool breed kept for both meat and milk production. It orginatedfrom local breeds of the Apennine crossed to Bergamasca (Ciani *et al*. 2014). Also in this case, despite of the crossbreeding with the Bergamasca, the breed has maintained a higher frequency of the G allele, corroborating again the hypothesis of the G allele being associated with hardness, and substantiating the conviction of the CD109 gene having a role in natural immunity.

CD109 is a protein-coding gene. Gene Ontology annotations related to this gene include serine-type endopeptidase inhibitor activity. To our knowledge, no studies were found in literature on the expression of this gene in livestock, however, in human medicine, this gene was associated to several types of cancer. CD109 gene expression was correlated with the pathological grade and tumor stage of some carcinomas (Hagikura *et al*.2010). Li *et al*. (2013) found that CD109 marker was expressed in the hepatic progenitor cells (HPCs) and suggested that this marker could potentially be utilized to identify and isolate HPCs for further cytotherapy of liver diseases. Finally, Bizet *et al*. (2011) identified CD109 as a TGF-β co-receptor and a negative regulator of TGF-β signaling. These authors demonstrated that CD109 increases binding of TGF-β to its receptors promoting localization of the TGF-β receptors into the caveolar compartment and facilitates TGF-β-receptor degradation. Thus, CD109 regulates TGF-β receptor endocytosis and degradation to inhibit TGF-β signaling.

The existing studies on immune response to infections, both in humans and in livestock, have compared the positive and the control groups using either the candidate gene approach, which is based on the analysis of pre-selected genes, or the GWAS, by exploiting anonymous marker panels of various density (Nalpas *et al*. 2013; Minozzi *et al*. 2010; Zare *et al*. 2014). However, so far, little overall consensus has emerged from these studies in terms of resistance loci, this being likely due, according to Riggio *et al*. (2014), to the apparent genetic complexity of the trait and the diversity of the studies, both for the considered breeds and the experimental approaches.

In the present study we proposed a novel application of the genotyping results obtained with panels of polymorphic markers; the new application consisting in uncovering the genes and the mutations that might play a direct role in the variation of the trait under investigation. Once the genes have been identified, specific q-RT PCR trials of gene expression for these genes could be designed, so to corroborate the effect of the markers. The recognition of genes that potentially play an important role in immune response in livestock will offer valuable reference in promoting sustainable animal farming.

## ACKNOWLEDGEMENTS

This study is part of the GENZOOT research program, funded by the Italian Ministry of Agriculture (Rome, Italy).

